# The rhythm of attentional stimulus selection during visual competition

**DOI:** 10.1101/105239

**Authors:** Sébastien M. Crouzet, Rufin VanRullen

## Abstract

Recent research indicates that attentional stimulus selection could in fact be a rhythmic process, operating as a sequence of successive cycles. When two items must be monitored, an intriguing corollary of this “blinking spotlight” notion could be that the successive cycles are directed alternately to each target; as a result, each item would effectively be selected at half the intrinsic rate of attentional selection. Here, we tested this prediction in two experiments. In an endogenous attention task, subjects covertly monitored one or two peripheral images in order to detect a brief contrast change. In the sustained occipital EEG power spectrum, selecting two vs. one item resulted in a relative increase around 4Hz and a relative decrease around 10–11Hz. In a second experiment, we tested if comparable oscillations could be observed in the stimulus-evoked EEG visual representational content. Subjects saw a first peripheral image displayed alone for 600ms, before a second one also appeared for the same duration, but at a different peripheral location. Using pattern analysis on EEG evoked-responses, we were able to create item selective classifiers that constantly indicated which stimulus was on the screen. The time-course of single-trial classifier decision values presented a relative spectral peak around 11Hz when only one object was present, and around 4–5Hz when two objects were on the screen. These results are both compatible with an attentional stimulus selection process sampling the visual field at around 10–11Hz, and resulting in a half-frequency effective sampling around 4–5Hz when there are two items to monitor.

## 1 INTRODUCTION

During natural vision, because of the limited processing capacity of the visual system, many different objects need to compete for representation. Visual attention regulates this competition by prioritizing specific objects at specific times. This is particularly emphasized in the biased competition account of attention (Desimone & Duncan, 1995; Desimone, 1998) in which attention resolves competition by biasing representations towards the attended object. Nevertheless, the simultaneous representation of multiple objects remains a challenge for any hierarchical system, including the current view of the ventral visual stream (Felleman & Van Essen, 1991; Riesenhuber & Poggio, 1999).

An elegant solution to overlapping representations of co-occurring objects is oscillatory multiplexing, whereby the representation of competing objects is segregated in time across different cycles or phases of one or more relevant oscillations (Bosman et al., 2012; Lisman & Jensen, 2013; Mclelland & Vanrullen, 2016). In multiplexing theories, attention is an oscillatory process with a rhythm corresponding to the α band, around 8 – 10 Hz (Busch & VanRullen, 2010; Fiebelkorn, Saalmann, & Kastner, 2013; Song, Meng, Chen, Zhou, & Luo, 2014; Landau, Schreyer, Van Pelt, & Fries, 2015). In such a scheme, attention drives the cyclic selection of co-occurring objects, represented one after the other through the alternating activations of each selective neural populations.

An intriguing hypothesis that can then be derived is that when two items are monitored, each item would effectively be selected/represented at half the intrinsic rate of attentional selection (or less than half, if additional time is required for attention to move in space). For example, Macdonald, Cavanagh, and VanRullen (2014) brought support for this hypothesis using reversal illusions in multiple simultaneously attended wagon-wheels, showing that the temporal-frequency tuning of the illusion decreased with the number of wheels to attend. Several recent behavioral studies used thorough designs and massive number of trials to indeed reveal fluctuations in behavioral performance itself (Landau & Fries, 2012; Fiebelkorn et al., 2013; Holcombe & Chen, 2013; Song et al., 2014). These results were compatible with (i) an intrinsic rhythm of attention and (ii) an under-sampling when attention is divided (Vanrullen, 2013). However, since the attentional cycle in these studies was defined based on a supposed reset by an external cue, their direct link with spontaneous brain fluctuations remains only hypothetical.

Using magnetoencephalography (MEG) and activity as a proxy for attention, Landau et al. (2015) brought more direct electrophysiological evidence that visual attention may oscillate around 4 Hz when subjects had to monitor two locations (which is half of an assumed 8 Hz intrinsic rhythm). Nonetheless, perhaps the most explicit demonstration came from single-cell recordings in the monkey inferotemporal (IT) cortex (Rollenhagen & Olson, 2005). They were able to show that (i) a subset of IT cells had an oscillatory spiking response to the onset of their preferred object, and (ii) these cells responded with the same 5 Hz periodicity when a second object appeared, but this time in anti-phase to their initial response. Such spiking behavior fits thoroughly with a multiplexing model using competitive interactions between neural populations selective to the two objects.

Here we designed two electroencephalography (EEG) experiments to test the half-sampling hypothesis when two objects are competing. First, in a spatial cueing task, we found that monitoring two vs. one item resulted in a relative increase of occipital EEG power around 4 Hz and a relative decrease around 10–11 Hz. In a second experiment, we tested if comparable oscillations could be observed in the stimulus-evoked EEG visual representational content using a protocol inspired by Rollenhagen and Olson (2005). Using time-resolved multivariate pattern analysis (MVPA), we were able to isolate a relative peak in decision value oscillations around 11 Hz when only one object was present, and around 4–5 Hz when two objects were on the screen. Both results are compatible with an attentional stimulus selection process sampling the visual field at around 10–11 Hz, and resulting in a half-frequency effective sampling (around 4–5 Hz) when there are two items to monitor.

## 2 MATERIALS AND METHODS

### 2.1 Participants

Thirteen (aged 21–32, 5 female) and nine (aged 21–33, 5 female) volunteers participated in Experiment 1 and 2 respectively. All subjects showed normal or corrected to normal eye function. Five participants were excluded from Exp1 because no attentional effect (facilitation of monitoring one vs. two locations) was found in their behavioral data (neither on reaction time nor on the contrast obtained from the adaptive procedure).

### 2.2 Stimuli and protocol

Stimuli were presented at a distance of 56 cm with a cathode ray monitor (resolution of 1280 × 960 pixels and refresh rate of 100 Hz) using the Psychophysics Toolbox (Brainard, 1997) running in MATLAB (MathWorks). The general stimuli layout was identical in both experiments: two images (or their noise counterpart) were presented on a gray background on the top-left and bottom-left (image angular size = 11.5^o^, horizontal and vertical eccentricity of respectively 7^o^ and 6^o^) of the display while observers held central fixation. This layout was preferred to left vs. right of fixation in order to increase the competition between the two stimuli by making them spatially close and within the same hemifield (Alvarez & Cavanagh, 2005). We assumed this would increase the likelihood to reveal an oscillatory process. The two images we used were two photographs, one face and one house, that had been shown previously to elicit very different neural representations in the ventral visual stream (Kriegeskorte et al., 2008). These images were first converted to grayscale, and their power spectra in the Fourier domain for each scale and orientation were averaged to cancel low-level differences. An additional noise image was created using this averaged power spectrum combined with the phase of random white noise. For a given participant the face and the house were both associated, when present, with a constant location throughout the experiment (e.g. upper-left for the house and lower-left for the face). This assignment was counterbalanced across participants. In Experiment 1, while holding central fixation, participants were instructed to monitor the “meaningful” image(s) (the face and/or the house) for the occurrence of a brief contrast increment (diameter = 1.7^o^, one-frame duration = 10 ms) in the center of one of the “meaningful” images (i.e. never on the phase-shuffled image location). Such target appeared on 50% of the 576 trials, at an unpredictable time (uniform random distribution between 1 and 7 s). The salience of the target (percent contrast change) was adapted using a QUEST procedure (Watson & Pelli, 1983) so that 80% were successfully detected on target present trials. We implemented a sliding version of the QUEST that took into account only the last 40 trials so that it would adapt to any change of threshold over the duration of the experiment. Once a target appeared, subjects had 1 s to push the space bar to respond before the trial ended. In case of a missed target, they received a negative feedback through a red “TARGET MISSED” message displayed for 1 s. Trials were separated by a pause of 2 s. In Experiment 2, after a random 400 to 800 ms fixation period, the two locations were filled with noise images, before one of the two images appeared at its assigned location (e.g. the face in the upper-left), followed by the other one (e.g. the house in the lower-left). Each step of this sequence lasted 600 ms. During the sequence, 0 to 3 targets (contrast increase of the image center as in Experiment 1) could appear at any location. After the sequence, a question mark replaced the fixation point, and subjects had to report the number of targets they perceived using the “v”, “b”, “n” and “m” keys (coding respectively for 0, 1, 2 and 3). This behavioral task was only there to maintain the involvement of participants in the experiment.

### 2.3 EEG recording and preprocessing

EEG and EOG were recorded at 1024 Hz using a Biosemi system (64 active and 3 ocular electrodes) and downsampled offline to 256 Hz. EEG data were filtered (pass-band from 1 to 40 Hz), re-referenced to average reference, epoched around trial onset (from -100 to +7000 ms in Exp1, from -200 to +1200 ms in Exp2), and baseline corrected (using the time period [-100 to 0] ms in Exp1 and [-200 to 0] ms in Exp2). The data were then screened manually to reject trials with noisy signal or typical artifacts (eye movements, blinks, muscle contractions, etc.): on average 15/576 trials were removed in Exp1 and 6.7/960 in Exp2. When appropriate, individual electrode data were interpolated by the mean of adjacent electrodes (on average 1.5 electrodes per subject were interpolated in Exp1 and 0.66 in Exp2).

### 2.4 Time-resolved multivariate pattern analysis (MVPA)

In Exp2, time-resolved multivariate pattern analysis (MVPA) was performed directly on the EEG electrode potentials. A decoding accuracy time-course was obtained by feeding the signal of all electrodes, independently for each time-point, to a linear classifier (L2-regularized logistic regression; Fan, Chang, & Hsieh, 2008). This method allows to find the optimal projections of the sensor space for discriminating between the two conditions (face first vs. house first). For each time point, the performance of the classifier was determined by using a Monte-Carlo cross-validation (CV) procedure (n = 120) in which the entire dataset was randomly split into a training set (80% of the trials) and a test set (the remaining 20% of the trials) at each CV. The splits were created to always contain an equal proportion of the two types of trial. On every CV, the signal of each electrode was normalized across trials (scaled between 0 and 1) by using parameters estimated from the training set only. The reported results of the MVPA procedure correspond to the average area-under-the-curve (AUC) across participants (and non-parametric 95% confidence intervals). For the subsequent frequency analysis, instead of AUC, we used the single-trial decision value time-courses so that potential oscillatory phase difference among trials would not be averaged out. These decision values correspond to the raw output of the logistic regression (before the thresholding used to produce the binary responses) passed through a sigmoid function to convert them to probability outputs.

### 2.5 Frequency analysis

In Exp1, trial durations could extend from 1 s to 7 s. Thus, the frequency analysis of the EEG signal was performed on overlapped segments (length of 750 ms, 80% of overlap) extracted from each trial. In Exp2, instead of the EEG signal, the frequency analysis was fed with the output of the MVPA analysis: the single-trial time-courses of the decision values. The two 600 ms segments corresponding to the first (one image) and second (two images) phase of each trial were used. Then, for both experiments, each segment (EEG signal or decision values) was z-scored, passed through an Hanning window to smooth the signal borders (the central 80% were kept intact), and zero-padded to span 2 s. The power of the frequency content of each segment was finally recovered using the squared magnitude of the Fourier series coefficients (based on the function fft() in MATLAB).

### 2.6 Frequency imbalance score (FIS)

In Exp1, to assess the behavioral impact of EEG frequency differences between conditions at the individual subject level, we computed an oscillatory frequency imbalance score (*FIS*). We first computed the frequency power difference (Δ*P*) between the conditions BOTH and ONE: more power in BOTH than ONE would result in a positive value, and vice-versa.

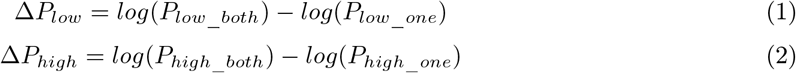

*P* corresponding to the power of the signal at a given frequency (low or high frequency clusters) and in a given condition (BOTH vs. ONE). According to our hypothesis, we should thus observe a positive value Δ*P_low_* associated with “low”-frequencies and a negative Δ*P_high_* associated with “high”-frequencies. Specifically, because there are 2 items to monitor, the “low” frequency range should be around half the “high” frequency range. At a second level, the frequency imbalance score (*FIS*) corresponded to the difference of these differences. The higher the *FIS*, the larger was the oscillatory power imbalance between the conditions BOTH and ONE.

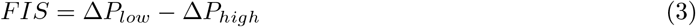

### 2.7 Statistical procedures

All error bars, except when otherwise specified, represent 95% bootstrapped confidence intervals of the mean across subjects. The contrast between conditions BOTH vs. ONE in Exp1 was tested for statistical significance using permutation tests in conjunction with cluster-based correction for multiple comparisons across frequencies (Maris & Oostenveld, 2007). Specifically, 2000 surrogate matrices were created by randomly permuting values between conditions across subjects. For each surrogate, a paired t-test between conditions was performed, p-values below a threshold of 0.05 were selected, and neighboring selected elements were aggregated into clusters. No specific constraint was set on the minimum number of marked elements in order to be considered a cluster. Elements were considered neighbors if they were directly adjacent in the frequency domain. Then, the t-values within each cluster were summed, and only the maximum cluster sum was saved for each surrogate. The distributions of these 2000 cluster-summed t-values reflected the null-hypothesis of no difference between the two conditions. Finally, the same procedure was performed on the original observed values and the true clusters considered significant only if their summed t-values was above the 95^th^ percentile of the surrogate null distributions. In Exp2, we restricted the analysis to the frequency range that appeared relevant based on Exp1 (from 2.5 Hz to 14 Hz) and used False Detection Rate (FDR) to control for multiple comparisons across frequencies.

## 3 RESULTS

### 3.1 Difference in EEG power spectrum between sustained monitoring of one vs. two items

In Experiment 1, we used an endogenous attention task with spatial cueing (Figure 1A) to compare EEG oscillatory power when subjects monitored one vs. two items. The locations to attend were indicated by the display of a meaningful image (e.g. a face, a house or both) instead of matched noise placeholders at the upper or lower left of the display. We chose to always present two stimuli (images or matched noise) on the screen to avoid any potential image low-level confound in the spectral measurements (on EEG spectrum in Exp1 and decoding spectrum in Exp2). A brief contrast change could appear on the monitored location at a random time from 1 to 7s after image onset. Participants had to press the space bar as soon as they detected this contrast change (target).

8 out of 13 participants had an expected facilitation of target detection when attending to ONE rather than BOTH locations, and were considered for further analysis (Figure 1D). Among these 8, half appeared to have a facilitation of detection accuracy, which manifested as a target contrast increase between the condition ONE and BOTH as selected by the QUEST procedure. For the other half, the effect of attention was seen on the reaction time distribution, with faster RT on average in the ONE than in the BOTH condition. Overall, the entire group of 8 subjects showed an average facilitation effect of 5% increase on target contrast (bootstrapped 95% CI across subjects = [3 11]%) and of 15 ms on RT (CI = [3 35] ms).

The key analysis focused on the EEG steady part of each trial: one second after image onset (i.e. attentional cueing, to leave time for attention to deploy and to ensure that visual evoked brain activity was not I ncluded) and before any target or motor response (Figure 1B and C, see Methods for further details). The frequency analysis of this steady-vision part of trials revealed two significant frequency clusters distinguishing the monitoring of one vs. two items (Figure 2A, see Methods for details about the statistical clustering procedure): a first positive (i.e. significantly more power in BOTH than ONE) low-frequency cluster ranging from 2.5 to 4.5 Hz (peak at 4 Hz), and a second negative (i.e. less power in BOTH than ONE) higher cluster at roughly twice the previous frequency ranging from 9.5 to 14 Hz (peak at 10.5 Hz).

**Figure 1:**
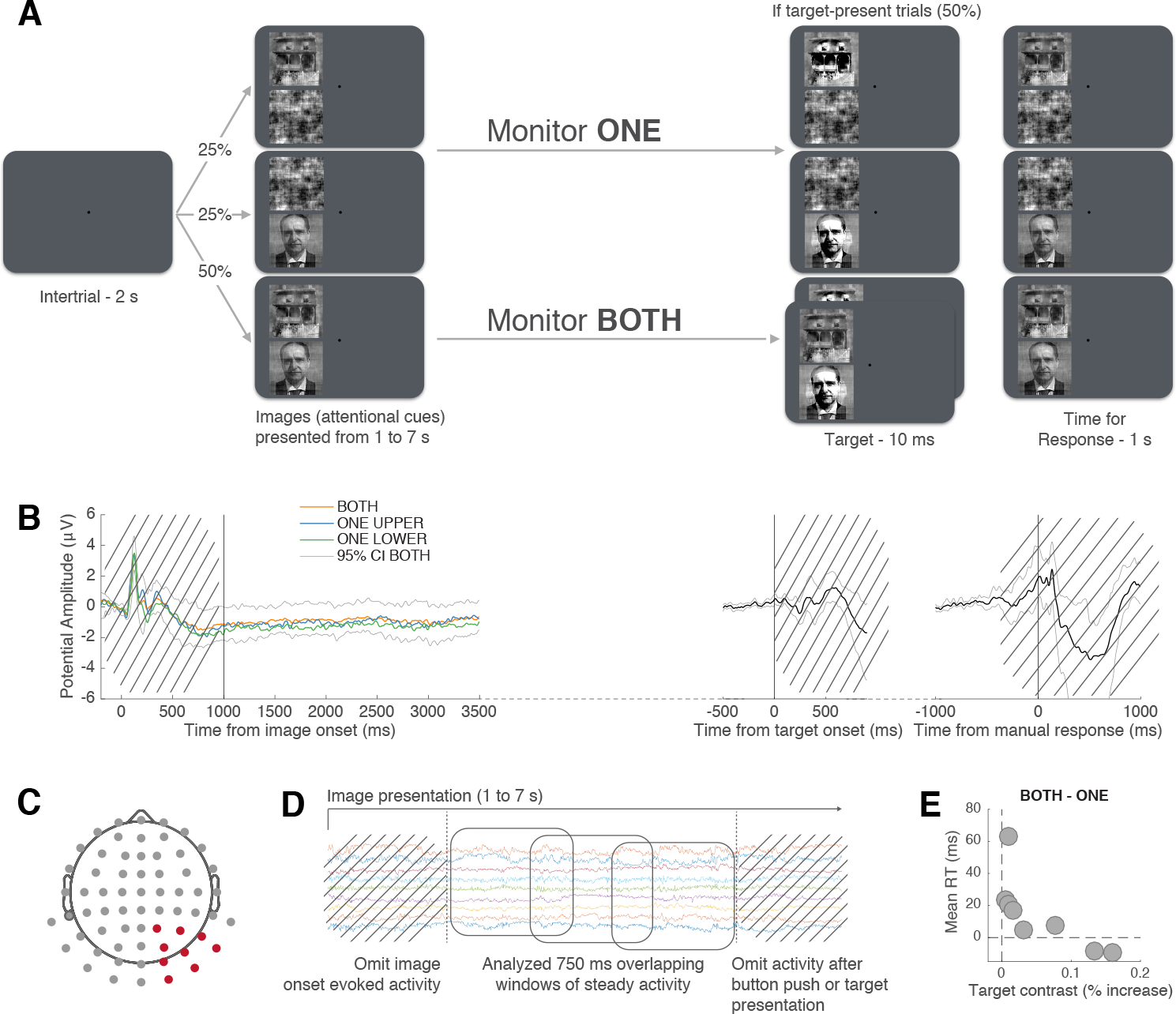
Experiment 1 methods, evoked activity and behavioral performance. (A) Protocol of Experiment 1. (B) Evoked Related Potentials (ERPs) locked to image onset (left), flash onset (considering only target present trials; middle) and motor response to the target (right). Main lines correspond to EEG activity time-course averaged across electrodes, trials and subjects, error bars to the bootstrapped 95% CI across subjects. Segments containing any such evoked responses were removed from subsequent frequency analyses (see Methods). (C) Because the task was visual and stimuli presentations were exclusively in the left hemifield, we selected and averaged together (only after performing the frequency analysis) the 9 right occipital electrodes. (D) Because trial durations were from variable lengths (1 to 7 seconds), the frequency analysis was performed on 750 ms overlapping windows (equivalent to three 4 Hz cycles) which were slided on the full steady activity found in each trial (see Methods for details). (E) Behavioral performance of the 8 subjects. Each point represents the difference between the conditions BOTH and ONE for each subject in terms of mean reaction time (RT) and average target contrast (selected by the adaptive procedure).

If these oscillations were associated with attention monitoring per se, then we should also observe an improved performance at target detection when they are present. Indeed, we found such a link (Figure 2B). First, the frequency imbalance score (FIS) between “low” and “high” frequency oscillations (= power difference [BOTH - ONE] at low-frequency minus power difference [BOTH - ONE] at high-frequency, see Methods) was inversely associated with reaction time (Figure 2B, repeated-measure two-way ANOVA: interaction between RT and frequency band; F(2,14) = 4.06; p = 0.04). In other words, the more powerful were the “low” and “high” oscillations, the faster were subjects’ responses. Looking independently at low (4–5 Hz) and high (10–11 Hz) oscillation clusters, it appears that each was in particular associated with the number of items to monitor: low and high-frequency oscillations modulating RT in the BOTH and the ONE condition respectively (Figure 2B). Notably, slow RTs were associated in both cases with close to zero power difference (between ONE and BOTH) in the corresponding frequency band.

Experiment 1 allowed us to demonstrate a switch in EEG power at specific frequency bands when subjects monitored one vs. two items. However, this correlation of oscillatory power (and behavioral performance) with the number of monitored items may only be indirectly linked with the information coded through these oscillations. Experiment 2 was designed to test this question more directly by examining the information contained in the EEG activity.

**Figure 2:**
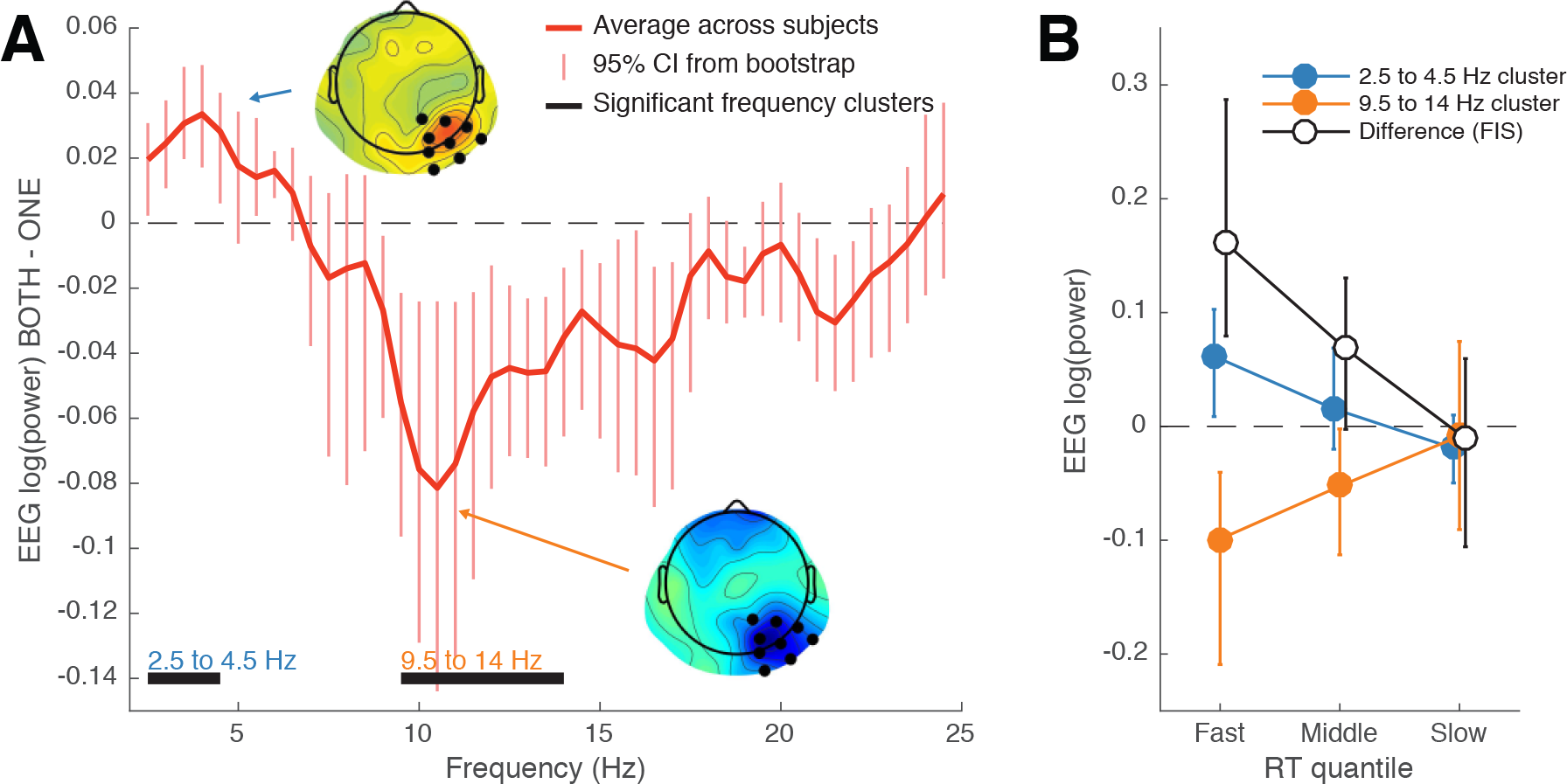
Results of Experiment 1. (A) EEG power spectrum difference between the conditions BOTH and ONE averaged across subjects (error bars correspond to bootstrapped 95% CI). The cluster-based permutation procedure revealed two significant clusters around 4–5 Hz and 10–11 Hz. Insets correspond to the topographies associated with each frequency cluster. (B) EEG power spectrum difference [BOTH - ONE], and Frequency Imbalance Score (FIS) as a function of reaction time quantiles. The FIS corresponds to the difference between the blue (“low” frequency cluster) and orange (“high” frequency cluster) lines. Not only did the imbalance vary with reaction times, but the link between oscillatory power and reaction time was specific to each frequency. More, a null difference in EEG power between conditions corresponded to the slowest reaction times.

### 3.2 Difference in decoding power spectrum between responses evoked by one vs. two objects

In Experiment 2, we designed an EEG experiment based on a protocol used previously in primates to investigate oscillatory responses to competing objects (Rollenhagen & Olson, 2005). This is a specific case where the visual system could take advantage of oscillations to represent multiple objects “simultaneously” (i.e. each object being associated to a different cycle of the support oscillations). Presenting two objects consecutively (one appears first and then the second is added to the display) to the monkeys on each trial, Rollenhagen and Olson (2005) showed that the firing rate of specific inferotemporal cells selective to one of the two objects (i) oscillates at 5 Hz in response to this object, (ii) appears to re-oscillate at 5 Hz, but in anti-phase to (i), when this object is presented second. This protocol seems to reveal oscillatory competition between populations of visual neurons. We designed an EEG experiment along these lines (Figure 3A): after fixation noise images appeared at the two placeholders (upper and lower left of the screen), then the first image appeared (e.g. face in the upper-left), and finally the second was also revealed (e.g. house in the lower-left). Each step of this trial sequence lasted 600 ms. Zero to three targets (a brief contrast change in one image) could appear during a trial. Participants had to report at the end of each trial how many they perceived.

The target counting task was only used in this experiment to make sure that participants were actively looking at the screen. Indeed, they reported the correct number of targets on 81.8% of the trials on average (see the confusion matrix in Figure 3B for detailed behavioral results). Depending on the number of targets presented, their performance was 96.7% (CI = [95.3 98.1]), 68.6% (CI = [65.7 74.3]), 80.8% (CI = [78.8 83]) and 81% (CI = [76.5 84.8]) for respectively 0, 1, 2 and 3 targets presented. Overall, participants thus had a high level of performance, attesting that they were properly involved in the experiment.

The aim of this experiment was to study the oscillatory component of the visual selective response to competing objects. As a substitute to the intracranial recording of object-selective cells used in Rollen-hagen and Olson (2005), we used classifiers trained on the activity of all EEG sensors. Training/testing a classifier at each timepoint independently allowed us to get a time-resolved index of the object-selective component of the EEG response. The use of such machine learning technique for the analysis of electrophysiological data in sensory experiments is now widespread (Ratcliff, Philiastides, & Sajda, 2009; Carlson, Tovar, Alink, & Kriegeskorte, 2013; Crouzet, Busch, & Ohla, 2015; Cauchoix, Crouzet, Fize, & Serre, 2016). The classification performance gives us an estimate of “how much” and “when” did the EEG activity differentiate the two conditions (face or house appearing first), reflecting the activation/deactivation of the face and house visual representations. The group-level results of this analysis are presented in Figure 3C: during the first part of the sequence (P1), the decoding time-course rises as early as 80/90 ms after image onset to reach a decoding performance (AUC) close to 0.8, and then slowly decreases after a few lower peaks. Overall, this dynamic is very similar to what is usually observed in visual experiments (see for example Carlson et al., 2013). The decoding from the second part of the sequence (P2) in which the second image is added to the display is less high (around 0.75) presumably due to the competition but still far above chance. Interestingly, during that second period the classifier is distinguishing between 2 conditions that are identical in terms of visual inputs (same 2 images at the same location on the screen) and only differ by their stimulation history (which of the 2 images appeared most recently).

**Figure 3:**
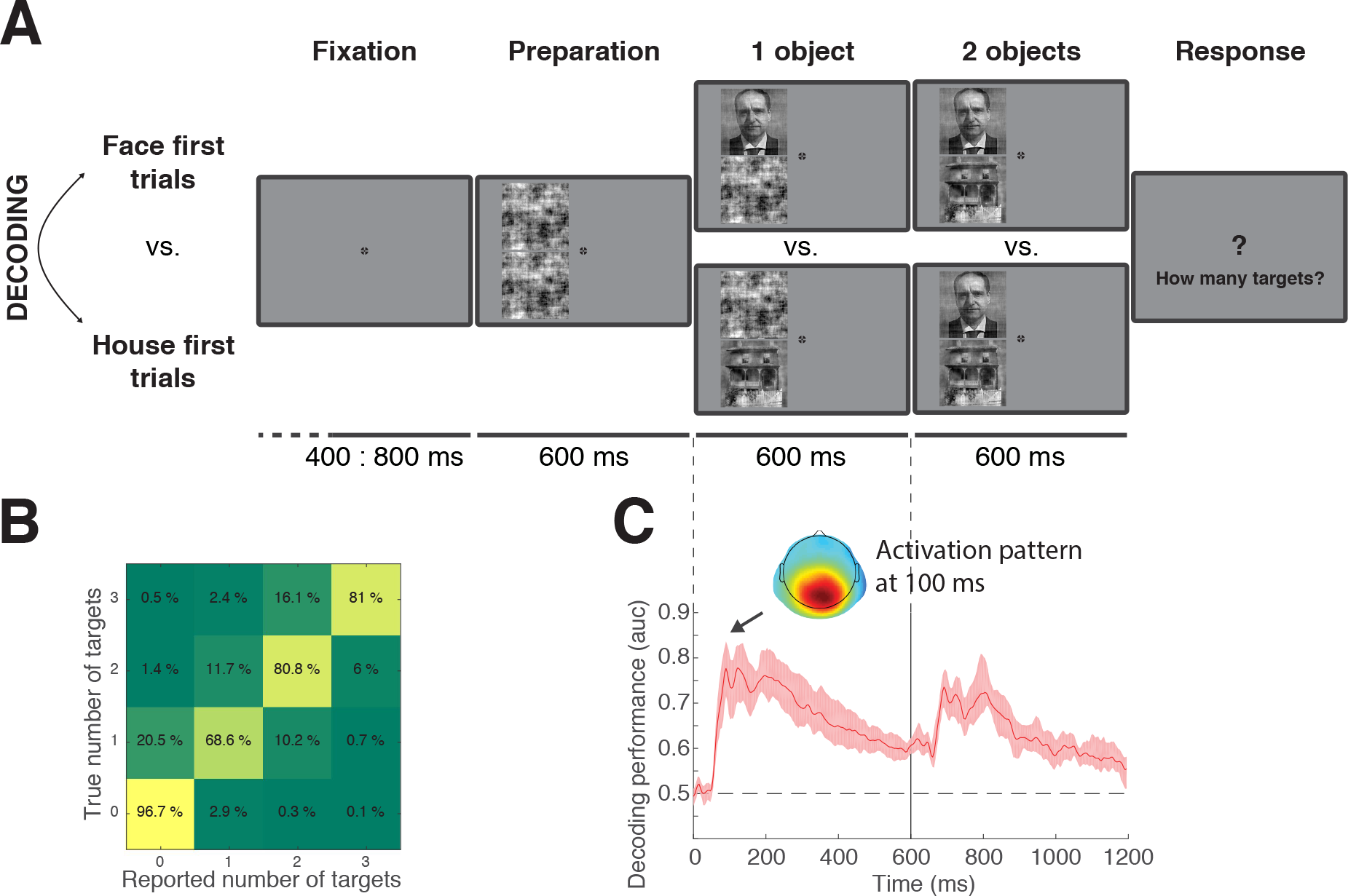
Experiment 2: protocol, behavioral and decoding performance. (A) Protocol of Experiment 2. (B) Confusion matrix between the number of targets presented and the number of targets reported. (C) Decoding performance time-course expressed in terms of AUC (area-under-the-curve) for the classification of the two types of trials (face first vs. house first). The activation pattern (illustration of the classifier weights) for the timepoint 100 ms after the first object onset appears as inset.

To measure the oscillatory content of the visual response associated with each period of the sequence (P1 and P2), rather than the cross-validated performance of the classifier (i.e. overall performance across trials as indicated by the AUC in Figure 3A), we used the cross-validated decision values obtained on each single-trial. When averaged, these decision values are by design equivalent to the AUC. However, considering them at the single-trial level allowed us to estimate and compare the oscillatory content of the classifier output in a way that is more robust to potential phase difference among trials. To that end, we averaged decision values obtained for a given trial across all the cross-validations in which this trial was in the test set, and performed the frequency analysis on each of these single-trial estimates (see Methods). The comparison of frequency distribution between P1 and P2 isolated two clusters of relative difference: more power in P2 than P1 around 4–5 Hz, and the opposite, less power in P2 than P1 around 10–11 Hz (Figure 4A and B). Individual subject results on these two selected clusters are presented in Figure 4C, showing a high consistency of the effect between participants. These results are both compatiblewith Experiment 1 as well as those of Rollenhagen and Olson (2005) in monkey single-cell recordings. Importantly, they support a model of multiplexed visual representations in which each object selected is represented at one cycle of a support oscillation fluctuating here at 10–11 Hz. Thus, when two objects are selected, they alternate and the effective sampling of each object becomes half of the original frequency, which indeed corresponds here to 4–5 Hz.

**Figure 4.**
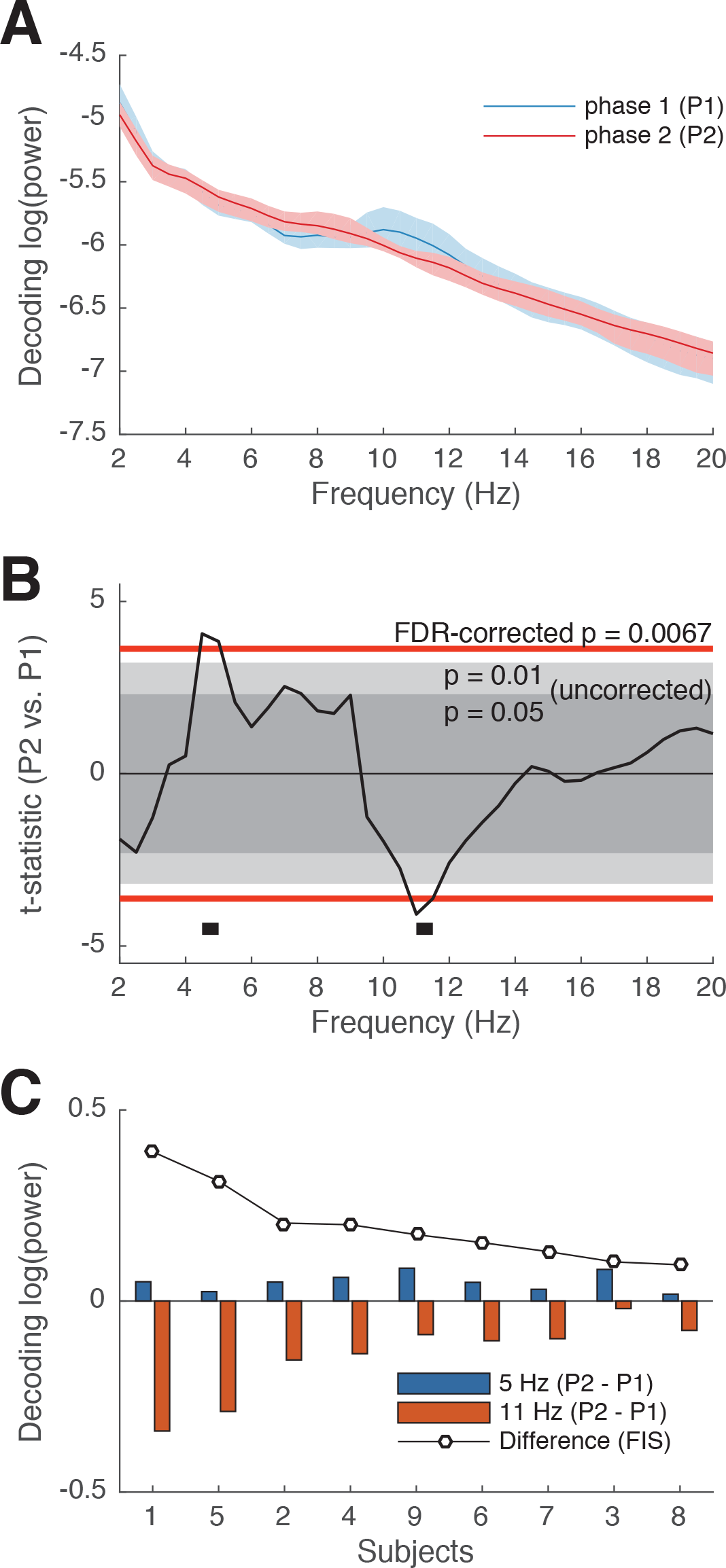
Results of Experiment 2. (A) Power spectra of the decision values associated with each part of the sequence. (B) Oscillatory power difference of the decoding decision values between the two parts of the trial sequence (t-statistics, red line corresponding to the FDR-corrected p-value). Two clusters reached significance: a low frequency one around 4–5 Hz and a high frequency one around 11 Hz. (C) Illustration of individual subject results: overall frequency imbalance (between low and high frequency for the difference between P2 and P1) for each subject and decomposed for the low and high frequency at the peak frequency of each cluster.

## 4 DISCUSSION

In two experiments, we observed variations of participants’ EEG oscillatory content depending on the involvement of visual competition. In Experiment 1, we observed a relative increase of subjects’ occipital EEG power around 4–5Hz and a relative decrease around 10–11 Hz when contrasting the monitoring of two vs. one locations for the appearance of a target. These variations of oscillatory power were associated with faster reaction times to detect the target in corresponding tasks: more power at 4–5 Hz associated with faster RT in the two-object task, more power at 10–11 Hz associated with faster RT in the one-object task. In Experiment 2, we tested if comparable oscillations could be observed in the stimulus-evoked EEG visual representational content. Using time-resolved MVPA, we observed a relative peak in decision value oscillations around 11Hz when only one object was present, and around 4–5Hz when two objects were on the screen. Overall, the specific frequency power modulations observed in both experiments were consistent with an attentional stimulus selection process sampling the visual field at around 10–11Hz, transfigured as a half-frequency effective sampling (around 4–5Hz) when there were two items to monitor.

If attention intrinsically samples the visual field at 10–11 Hz, the brain regions producing this intrinsic rhythm of attention should show the same activity power spectrum regardless of the number of attended locations. How can we observe a half-rhythm in the EEG frequency spectrum when subjects are monitoring two objects? The fact that we are able to observe these types of modulations in our paradigm likely implies that what we are recording is not the source of rhythmic attentional sampling (originating from attentional control areas) whose frequency would remain constant no matter the number of monitored object(s). Instead, what we are observing here is likely to be the result of attentional modulation of sensory processing (i.e. how attention modulates object representations in sensory areas). Indeed, if we assume that what we record at the level of a single electrode activity is not the source but the result of attentional selection, then even if the source of attentional modulation oscillates at 10 Hz independently of the condition, we can expect the activity of each electrode to be biased towards one of the objects/locations to represent (i.e. responding more to one than the other). In that case, the fluctuations recorded at this specific electrode would be associated with the attentional modulation of the object representation itself. In other words, if one object is selected once every two cycles, then it will be “attended at 4–5 Hz”, and any electrode with a response biased toward this object would show a power increase around 4–5 Hz.

A related question is by how much we expect the intrinsic rhythm to be divided when the observer is attending multiple locations instead of one. We will consider here the case of our experiment where there are two locations to monitor. In an idealized system in which there is no switching cost, the ratio should be exactly two: an intrinsic rhythm of 10 Hz would result in a half-rhythm of 5 Hz. However, it is reasonable to consider that additional processes/time/machinery might be needed when the spotlight switches between locations, rather than staying in place on each successive sample. This additional time would result in a division factor that would be slightly above 2, as found here. Another (not exclusive) possibility is that the attentional spotlight is naturally linked to the position of the eyes, and thus sometimes returns to the central fixation point between successive “attentional” samples of the targets. In the extreme version of this, the monitoring of two target locations would effectively imply 3 locations to sample, and be reflected in the EEG by a division factor of 3. Our results, with a dividing factor slightly above 2 (ratio between 11 Hz and 4.5 Hz is 2.44), could be compatible with various combinations of these explanations, and further research will be needed to properly answer this question.

Overall, our results directly support the hypothesis that the representations of competing objects alternate over time. They are thus compatible with oscillatory multiplexing models such as those proposed by Lisman and Jensen (2013) and Fries (2009) (see also Mclelland & Vanrullen, 2016), as well as consistent with many other experimental findings (Busch & VanRullen, 2010; Landau & Fries, 2012; Landau et al., 2015). But beyond being consistent with a general class of oscillatory models of perception, how do our findings relate to other experimental studies in terms of the specific frequency bands involved? Various intrinsic perceptual/attentional rhythms have been reported in the literature: around 7–8 Hz (Busch, Dubois, & VanRullen, 2009; Busch & VanRullen, 2010; Landau & Fries, 2012; Fiebelkorn et al., 2013; Vanrullen, 2013; Landau et al., 2015; Dugué, McLelland, Lajous, & VanRullen, 2015), 10 Hz (Mathewson, Gratton, Fabiani, Beck, & Ro, 2009;Dugué, Marque, & VanRullen, 2011; Dugué & VanRullen, 2014; Macdonald et al., 2014) or 13 Hz (VanRullen, 2006; Drewes & VanRullen, 2011), with corresponding half-rhythms that would thus range from 3.5 to 6.5 Hz. The results of our two studies are in the middle range of this spectrum: we found intrinsic and divided rhythms at 10–11 Hz and 4–5 Hz respectively. Could these variations in the frequency range reported across studies actually correspond to functional differences? A dichotomy was recently suggested between rhythmic processes that would be more attentional or more perceptual in nature (VanRullen, 2016): attentional processes would operate under an intrinsic 7–8 Hz rhythm (Busch et al., 2009; Busch & VanRullen, 2010; Landau & Fries, 2012; Fiebelkorn et al., 2013; Vanrullen, 2013; Landau et al., 2015; Dugué et al., 2015) while more perceptual processes would rather operate at around 10 Hz (Dugué et al., 2011; Dugué & VanRullen, 2014; Macdonald et al., 2014). Our results are in apparent contradiction with this suggestion, since our task involves spatial attention but the intrinsic rhythm is around 10–11 Hz. However, many other factors could also determine frequency variations: the response modality (e.g. manual vs. saccadic), the eccentricity or contrast of the stimuli, the distance between locations to monitor, or even the overall task difficulty. Future work may help to characterize the relation between rhythmic frequency and these various factors.

